# The Influence of Spatial Frequencies, Orientation and Familiarity on Face Stimuli Integration

**DOI:** 10.1101/2025.02.11.637641

**Authors:** Maria Cuomo, Giovanni Federico, Alessio Fracasso, Antimo Buonocore

## Abstract

When we observe an object, our visual system identifies its shape and integrates it with specific details to form a coherent representation. This *coarse-to-fine* approach involves rapid processing of low spatial frequency (LSF) content to generate a basic template, which aids the integration of the more detailed high spatial frequency (HSF) information. Here we explore with two experiments how the contribution of LSF and HSF integration extends to face processing. To do so, we leveraged the face inversion effect, whereby inverted faces are more difficult to recognize than upright ones. In Experiment 1, ten participants matched two familiar faces displayed in rapid succession (template and probe face, respectively). The template and the probe shared either the same SF (congruent) or had complementary SF (incongruent). In congruent conditions, HSF templates yielded better matching accuracy than LSF templates. However, in incongruent conditions, mapping LSF probes onto HSF templates was more effective, but only for upright faces. We propose that, depending on the task, holistic processing may be facilitated by detailed information. In Experiment 2, twelve participants performed the same task with both familiar and unfamiliar faces. While for familiar faces the effects were the same as Experiment 1, for unfamiliar faces the overall accuracy was better for congruent than incongruent conditions, and, crucially, it was independent of the template SF. Our results challenge the view that LSF content provides a foundational template for integrating HSF information, and instead suggest a flexible encoding of SF information, that depends on image contingencies.

## Introduction

Natural environments consist of a multitude of complex objects. Yet, humans can categorize an entire scene or specific items in less than 150 milliseconds (Grill-Spector & Kanwisher, 2003; Potter & Levy, 1969; Thorpe et al., 1996). This rapid processing demonstrates the exceptional efficiency of visual perception and recognition with minimal cognitive effort (DiCarlo et al., 2012; Groen et al., 2017). A substantial body of research emphasizes the critical role of spatial frequency content in image processing and categorization (Collin, 2004; Guyader et al., 2004; Hughes et al., 1996), proposing that visual stimuli are initially decomposed into fundamental components based on their spatial frequency spectrum (DeValois & DeValois, 1991), which mirrors the hierarchical organization of the visual system. At the neurophysiological level, spatial frequencies are in fact processed through distinct pathways, starting from the retinal output and extending to the visual cortex, reflecting anatomical and functional specializations within the visual system (Livingstone & Hubel, 1988). Low spatial frequencies (LSFs), conveyed via the faster magnocellular pathway (Felleman & Essen, 1991; Livingstone & Hubel, 1987), provide a global or coarse representation of the scene, enabling rapid identification of general shapes and spatial layouts. In contrast, high spatial frequencies (HSFs), which are transmitted more slowly through the parvocellular pathway, support the perception of finer details, such as edges and borders (Bar et al., 2006; Kauffmann et al., 2014). This information is further relayed through the dorsal stream, where LSF are predominant. On the other hand, HSF information conveying fine details, moves predominately along the ventral stream (Skottun, 2015).

According to the neurophysiological description, experiments in primates (Bullier, 2001) and psychophysical studies in humans (Ginsburg & Arthur, 1980; Hughes et al., 1996) propose that spatial frequency processing typically follows a coarse-to-fine sequence of signal integration. First, the visual system identifies an object’s general shape using LSF, then, it progressively integrates finer details conveyed via HSF to construct a coherent representation (Bar et al., 2006; Bullier, 2001). In the predictive coding framework (Rao & Ballard, 1999), Bar et al. (2006) proposed that LSF are quickly relayed to the orbitofrontal cortex to generate predictions, which are fed back to inferotemporal areas to guide HSF processing and enable rapid recognition (Bar et al., 2006; Kveraga et al., 2007). This is reflected in differences in processing time, as demonstrated by numerous studies using simple stimuli with low-level characteristics, such as gratings. In support of this view, research consistently shows that perceiving HSF requires longer response times compared to LSF (Breitmeyer & Julesz, 1975; Broggin et al., 2012; Vassilev et al., 2002). Additionally, electrophysiological studies in humans have shown that early components of the visual evoked potential, such as the P1 wave, exhibit larger amplitudes in response to LSF than to HSF (Proverbio et al., 1996). Similarly, in primates, both visual (e.g., V1) and visuo-motor (e.g., SC) areas exhibit stronger and faster responses to LSF stimuli (Chen et al., 2018). These neurophysiological findings align with behavioral observations in humans, showing that complex natural stimuli, such as objects and faces —primarily composed of low- and mid-spatial frequencies, play a significant role in attentional orienting, eye movement programming (Bogadhi et al., 2020; Buonocore et al., 2020; Fracasso et al., 2015), and inhibitory selection mechanisms (Buonocore & Hafed, 2023; Taylor et al., 2024).

Unlike gratings and other non-face simple objects, faces exhibit a higher degree of complexity, requiring the integration of both internal and external features (e.g., eyes, nose, mouth, hair) to create a coherent percept (Goffaux & Rossion, 2006). This process, known as holistic face processing, involves combining facial features into a unified perceptual whole rather than analyzing them independently (Sergent, 1984; Tanaka & Farah, 1993; Young et al., 1987). For example, Sergent and Hellige (Sergent & Hellige, 1986) proposed that holistic face processing utilizes distinct regions of the spatial frequency spectrum. Specifically, LSF convey holistic information, such as pigmentation and coarse shading cues, while HSF transmit finer featural details, such as the eyes or mouth (Goffaux & Rossion, 2006; Maurer et al., 2002). Interestingly, also face processing seems to follow a coarse-to-fine integration process, with LSF (global shape) processed faster than HSF (internal details) (Goffaux et al., 2011; Halit et al., 2006; Hegdé, 2008; Nakashima et al., 2008; Petras et al., 2019; Schuurmans et al., 2023). These predictive mechanisms remain active across eye movements, as peripheral information about face stimuli is subsequently foveated (Buonocore et al., 2020; Huber-Huber et al., 2019).

Several behavioral effects demonstrate that faces are processed holistically: the part-whole effect (Tanaka & Farah, 1993), the composite effect (Young et al., 1987) and inversion effect (Yin, 1969). In particular, the inversion effect shows how face recognition is disproportionately impaired by inversion compared to the recognition of other objects (Rossion & Gauthier, 2002). Additionally, this effect is characterized by a delayed and larger N170 in response to inverted faces (Eimer, 2000; Rebai et al., 2001; Rossion & Gauthier, 2002). Hence, face inversion effect has been interpreted as indicating disruptions in either configural processing (the spatial relationships between facial features) or holistic processing (Itier & Taylor, 2002; Maurer et al., 2002; Rossion, 2008; Rossion & Gauthier, 2002; Tanaka & Farah, 1993; Taubert et al., 2011). Interestingly, some evidence suggests that face inversion has a more pronounced effect on LSF processing (Collishaw & Hole, 2001; Nagayama et al., 1995; Williams et al., 2009). For example, Hsiao (Hsiao et al., 2005) showed that the inversion effect primarily impacted LSF faces, delaying the M170 but did not significantly affect HSF faces. The author interpreted this result suggesting that LSF might be processed more like objects. These findings stand in contrast to earlier research, which posits that the face inversion effect is not dependent from spatial frequency processing (Collin, 2004; Collin et al., 2004; Gaspar et al., 2008; Willenbockel et al., 2010).

Low-level features are not the only factors that influence the representation of stimuli. In daily life, we are constantly surrounded by numerous faces, ranging from unfamiliar individuals to acquaintances, loved ones, and well-known celebrities. Extensive research shows that the perception of familiar and unfamiliar faces differs in several significant ways (Johnston & Edmonds, 2009; Young & Burton, 2018). Recognizing familiar faces is easier and more accurate than identifying unfamiliar ones (Ellis et al., 1978; Klatzky & Forrest, 1984), since familiar face recognition might rely more on relation between internal features, while recognition of unfamiliar faces might be related to featural information (Vereš-Injac & Persike, 2009). On a similar note, a study by Castello and colleagues (Castello, Wheeler, et al., 2017) investigated holistic and feature-based face processing under face inversion for both personally familiar and unfamiliar faces. The findings revealed an advantage for familiar faces in both upright and inverted conditions, because the familiarity facilitates feature-based face processing. Although the effects of familiarity and face inversion in face recognition are well documented in the literature, it is currently not clear whether and how they might interact with the low-level visual features of the presented stimuli.

In the present study, we measured performance in a face recognition task in which we varied the spatial frequency, the orientation and the familiarity of our stimuli. In agreement with previous literature, upright faces should promote holistic processing, characterized by an integrated, global analysis of spatial frequencies, whereas inverted faces should encourage feature-matching strategies that emphasize local, specific elements of facial information (Itier & Taylor, 2002; Rossion & Gauthier, 2002; Schwaninger & Mast, 2005; Tanaka & Farah, 1993; Taubert et al., 2011). If face processing strictly follows a coarse-to-fine mechanism, we predict that recognition performance will be enhanced when HSF probes are matched with LSF templates, regardless of orientation. Alternatively, task demands required for stimulus normalization (e.g., mental rotation) may modulate this integration process differently, potentially leading to task-dependent variations in how spatial frequencies are utilized and combined for face matching (Schyns, 1998; Schyns & Oliva, 1996). These variations may also depend on stimulus familiarity. To empirically test these hypotheses, we designed two experiments. In the first experiment, participants completed a match-to-sample task in which familiar face templates, either low-pass or high-pass filtered, were followed by a second face probe presented either upright or inverted. Experiment 2 extended this design by introducing unfamiliar faces alongside familiar ones to examine the role of familiarity. In both experiments our results diverge from classic coarse-to fine models (Bar et al., 2006) and indicate that for upright stimuli, matching performance is enhanced when LSF probes are matched to HSF templates. This result suggests an integration process in which fine information (HSF) supports the interpretation of coarse information (LSF), likely by providing more accurate templates for the representation of the face. In contrast, inverted faces, which disrupt holistic processing, promoted feature-matching strategies and lead to better performance when HSF probes, were matched with LSF templates. In Experiment 2, spatial frequencies of the templates did not modulate the perception of unfamiliar faces.

Taken together, these findings reveal that the integration of spatial frequencies operates differently based on face orientation and familiarity. They emphasize the critical role of orientation in modulating how spatial frequency information is utilized during face recognition (Castello, Wheeler, et al., 2017; Goffaux & Rossion, 2006).

## Experiment 1: The effect of spatial frequency and orientation on face perception

### Method

#### Participants

A convenience sample of ten participants (9 females and 1 male; mean age 24 years, SD = 1.5) was recruited to participate in the study. All participants self-reported no history of neurological or visual impairments and had normal or corrected-to-normal vision. They were naïve to the purposes of the experiment and primarily drawn from university students and staff who participated voluntarily. Written informed consent was obtained, in accordance with the 1964 Declaration of Helsinki. Ethical approval was granted by the local ethics committee at the college of Medical, Veterinary and Life Sciences, University of Glasgow.

#### Apparatus and stimuli

Stimuli were generated using MATLAB R2021a (MathWorks, Inc.) and Psychophysics Toolbox 3.0.18 (Brainard 1997; Pelli, 1997). All the stimuli were in gray scale on a gray background (RGB = 126), presented on a 27-inch LCD monitor with a resolution of 1920 by 1080 pixels and a refresh rate of 144 Hz (6.9 ms). The participants’ eyes were aligned with the center of the display at a distance of 65 cm. The fixation point was a small black (RGB = [0, 0, 0]) dot of about 0.30 (10 px) degrees radius. All the stimuli were presented at the center of the screen. As perceptual stimuli, black and white images (about 7.76 by 7.76 degrees of visual angle) (256 x 256 px) of five male and five female celebrity faces were used in Experiment 1. These faces were selected from the Celebrities in Frontal-Profile (CFP) dataset (Sengupta et al., 2016), which contains images of 500 different celebrities. Each celebrity is represented by 10 frontal images and 4 profile images. In this study, we exclusively selected frontal faces with neutral expressions to minimize the influence of pose and expression on the results. We then created two datasets of stimuli, depending on the type of filter applied. For the first set of images, we applied a low pass Gaussian filter under 1 cycle per degree to the face images. For the second set of images, we applied a high pass Gaussian filter above 1 cycle per degree. The mask was a square of about 7.76 by 7.76 degrees of visual angle made of random pixels with a random level of gray between RGB equal to zero (black) and 255 (white), centered on the template stimulus. After applying the mask, the luminance and spatial frequency content of the face images were homogenized throughout using the SHINE toolbox (Willenbockel, Sadr, et al., 2010). Afterward, we cropped the images into a circle, centered at the middle of the image with a radius of about 3.03 degrees of visual angle (100 px), in order to exclude facial features such as ears and hair. Participants were tested in a dark room, except for the display monitor, and the operator monitor located behind the participant and facing away from them.

#### Procedure

Participants were positioned on a chin-and-forehead support to ensure head stability. On each trial, participants were instructed to fixate a dot presented at the center of the screen. Following a variable delay of 1000 to 1500 ms after the onset of the fixation dot, an upright face stimulus (*template*) was presented for 200 ms, followed by a mask displayed for 100 ms (Fig. 1A). The template could be filtered either with a low pass or high pass filter (see Apparatus and stimuli). Following the presentation of the mask, there was an additional 200 ms blank interval after which a second face (*probe*) was presented for 200 ms. Depending on the condition, the probe could have been either the same or a different identity of the template. Critically, the probe could be filtered with a low pass or high pass filter. Finally, the probe could be oriented either upright or inverted (180-degree rotation) (Fig. 1B). Participants were required to determine whether the identity of the second face (the probe), was the same as or different from that of the first face, the template (two-alternative forced choice, 2AFC). The gender identity of the probe was always the same as the template.

**Figure 1.**
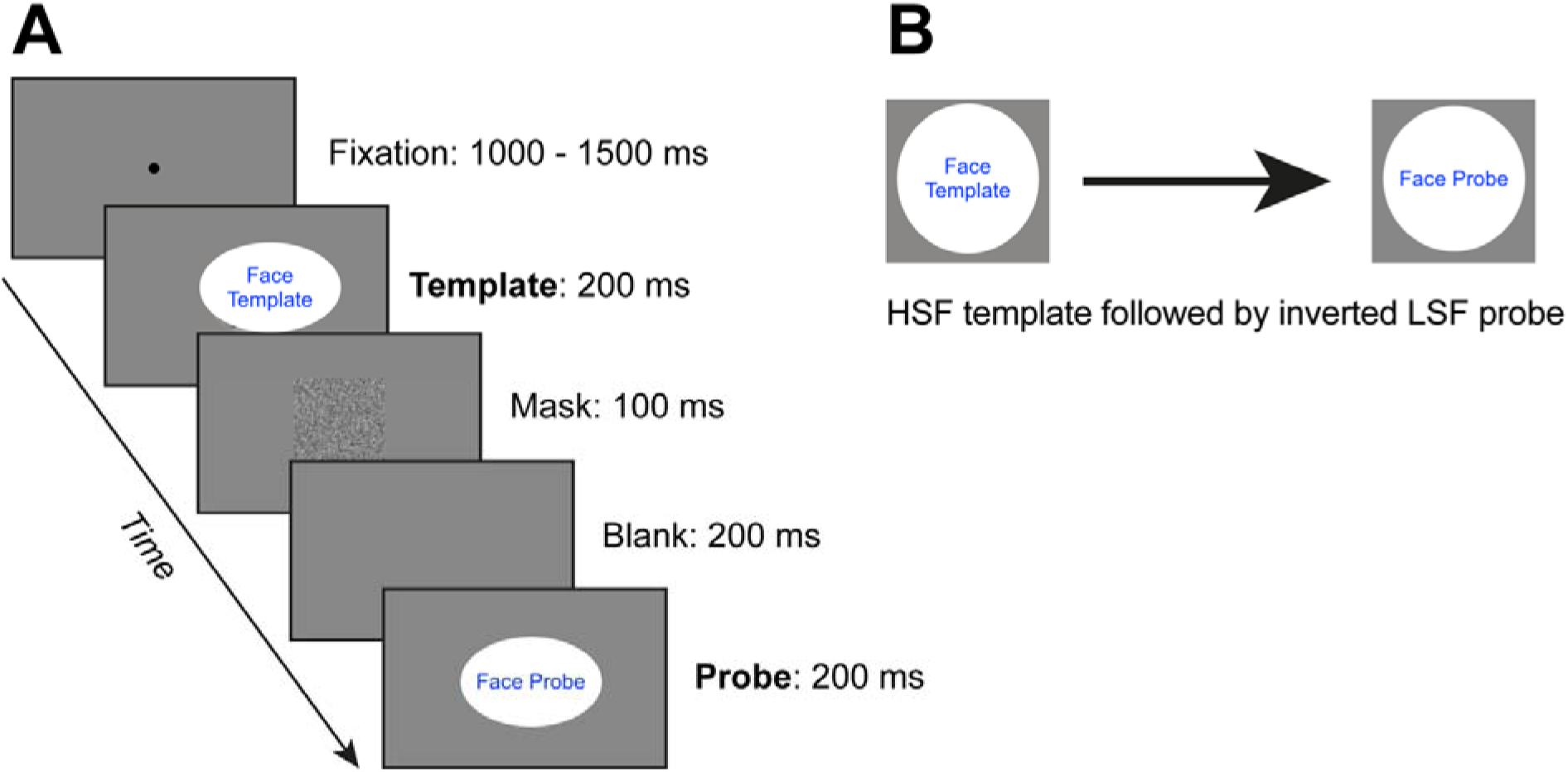
Procedure and stimuli used in Experiments 1 and 2. **(A)** Participants fixated on a central dot for a variable duration between 1000 and 1500 ms. Then, a face stimulus (template) was presented for 200 ms, followed by a mask (100 ms) and a blank screen (200 ms). After the blank screen, a second face stimulus (probe) appeared for 200 ms. The probe could either match or differ from the template in identity and spatial frequency. Additionally, the probe was presented either upright or inverted. **(B)** The diagram illustrates a condition in which a high-spatial-frequency (HSF) template was followed by a low-spatial-frequency (LSF) inverted probe. In Experiment 1, all face stimuli were selected from the Celebrities in Frontal-Profile (CFP) dataset (Sengupta et al., 2016). In Experiment 2, a new set of unfamiliar faces was selected from the dataset by Willenbockel et al. (2010).

Responses were recorded on a standard keyboard by pressing the ‘a’ key for the same identity and the ‘d’ key for different identities. There was no time limit for responses and after each response, the next trial began. After each block of trials (see Experimental design below), participants could take a short break. They started the next block of trials by pressing any key on the keyboard. At the end of the entire session all participants were debriefed about the research aim. Each session lasted about 40 minutes, breaks included.

#### Experimental design and statistical analysis

The full design comprised 16 experimental conditions, determined by the factorial combination of four factors: *Validity* (validity between template and probe: valid vs. invalid), *SF Congruency* (the congruency between the spatial frequency of the template and the probe: congruent vs. incongruent), *Template SF* (the spatial frequency of the template: LSF vs. HSF), and *Orientation* (upright vs. inverted). For the statistical analysis, data were aggregated across the Validity factor. By convention, the naming of the single conditions followed the ordering of the factors; for example, a low spatial frequency template (LSF) followed by high spatial frequency probe (HSF) (incongruent condition) in the inverted condition (i) would be named LSF-HSFi. Accordingly, all the conditions were named as follows: LSF-LSFu, LSF-LSFi, HSF-HSFu, HSF-HSFi, LSF-HSFu, LSF-HSFi, HSF-LSFu, and HSF-LSFi. Each of the eight conditions was repeated 40 times. Participants performed a total of 320 trials divided in 10 blocks of 32 trials each. We collected a total of 3200 trials across all participants in Experiment 1.

Subsequently, for each participant and condition (SF Congruency by Template SF by Orientation), we calculated mean accuracy and reaction times (RTs). Mean RTs were calculated only for correct trials. Due to a technical problem, RTs were not recorded for the first two participants. The standard error of the mean (SEM) was also calculated for each of the means described above. The effects of *SF Congruency* (congruent vs. incongruent), *Template SF* (LSF vs. HSF), and *Orientation* (upright vs. inverted) on accuracy (0, 1) and reaction times were analyzed using mixed-effects logistic regression (GLME) using “simple coding” to determine the main effects and the interactions. The models are summarized in Eq. 1 and Eq. 2 in Wilkinson notation (Wilkinson and Rogers, 1973):

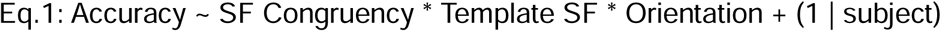

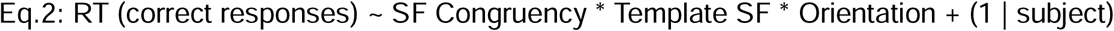

To further investigate the three-way interaction between SF Congruency, Template SF, and Orientation, the data was first divided into the two SF Congruency conditions (congruent vs. incongruent). Then, for each subset of data, a GLME on accuracy (0, 1) was conducted, examining the effects of Template SF and Orientation. The models are summarized in Eq. 3.1 and Eq. 3.2:

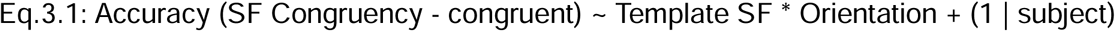

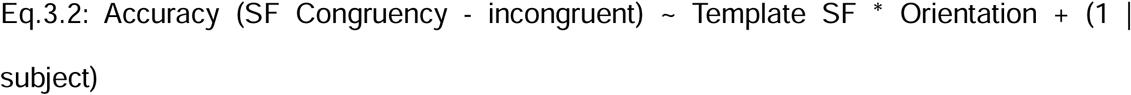

For the GLME, follow-up comparisons among pairs of means were further tested with an F statistic via specification of contrast matrices for the critical conditions. For the repeated measure ANOVA, we used pairwise t-test comparisons. The alpha level was set at 0.05 and corrected for multiple comparisons (Bonferroni correction). All data pre-processing and statistical analyses were conducted with custom scripts in MATLAB R2019a (MathWorks, Inc.).

## Results

### Face matching is influenced by both template and probe features

We first investigated whether the perception of the probe was affected by its orientation and spatial frequency relative to the previously presented template. Overall accuracy in the probe familiar matching task, was generally high (87% correct) and varied across experimental conditions. The GLME analysis (Eq. 1, see Table 1) revealed main effects of SF Congruency (β = 1.1772, 95% CI [0.65722, 1.6972], t = 4.4391, p = 9.38*10), indicating that congruent trials, in which participants were exposed to the same probe as the template, led to higher performance compared to incongruent trials (congruent: M = 0.91, SEM = 0.02; incongruent: M = 0.83, SEM = 0.03) (Williams et al., 2009). A main effect of Template SF (β = 0.66455, 95% CI [0.21112, 1.118], t = 2.8738, p = 0.004086) showed that participants were generally more accurate with HSF templates (M = 0.88, SEM = 0.02) compared to LSF templates (M = 0.86, SEM = 0.03).

**Table 1.**
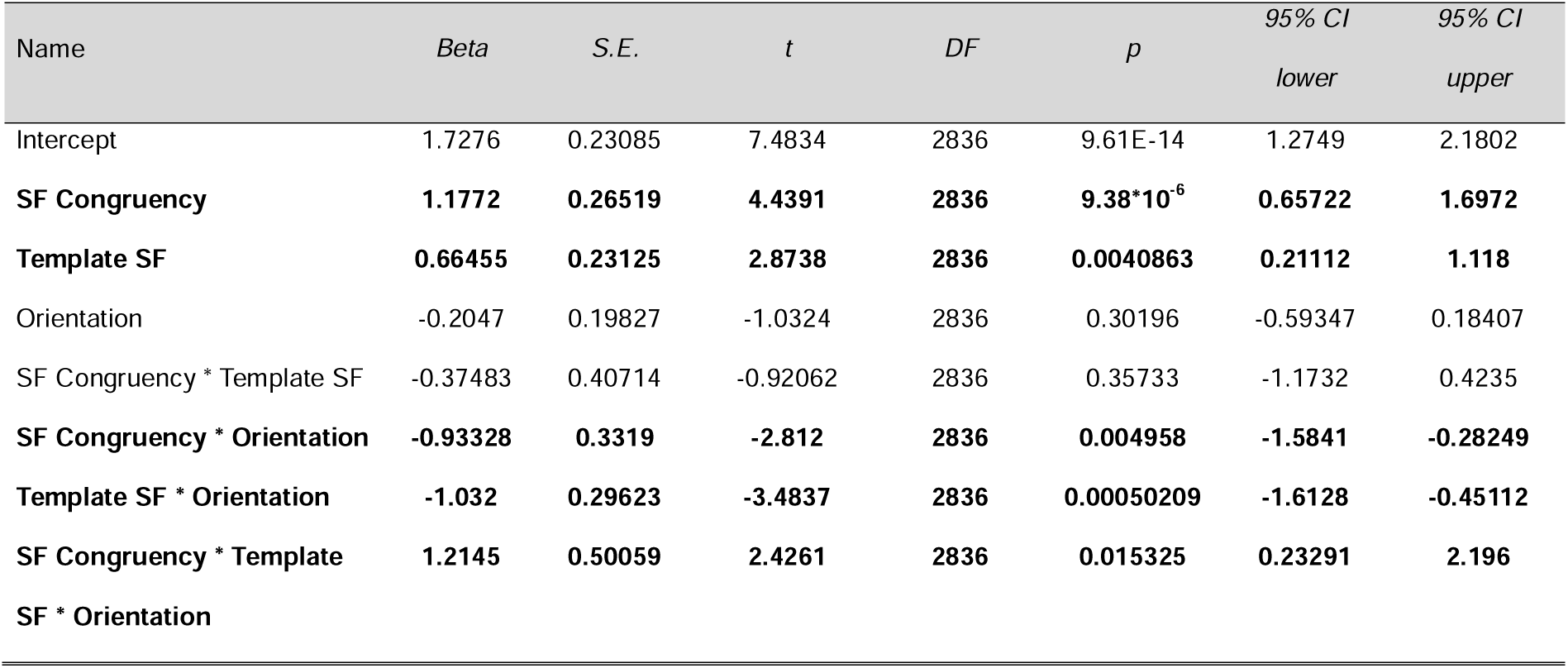
Experiment 1 GLME parameters for accuracy. Significant parameters are depicted in bold.

| Name | *Beta* | *S.E.* | *t* | *DF* | *p* | 95% *CI*<br>*lower* | 95% *CI*<br>*upper* |
| --- | --- | --- | --- | --- | --- | --- | --- |
| Intercept | 1.7276 | 0.23085 | 7.4834 | 2836 | 9.61E-14 | 1.2749 | 2.1802 |
| **SF Congruency** | **1.1772** | **0.26519** | **4.4391** | **2836** | **9.38\*10<sup>-6</sup>** | **0.65722** | **1.6972** |
| **Template SF** | **0.66455** | **0.23125** | **2.8738** | **2836** | **0.0040863** | **0.21112** | **1.118** |
| Orientation | -0.2047 | 0.19827 | -1.0324 | 2836 | 0.30196 | -0.59347 | 0.18407 |
| SF Congruency * Template SF | -0.37483 | 0.40714 | -0.92062 | 2836 | 0.35733 | -1.1732 | 0.4235 |
| **SF Congruency \* Orientation** | **-0.93328** | **0.3319** | **-2.812** | **2836** | **0.004958** | **-1.5841** | **-0.28249** |
| **Template SF \* Orientation** | **-1.032** | **0.29623** | **-3.4837** | **2836** | **0.00050209** | **-1.6128** | **-0.45112** |
| **SF Congruency \* Template SF \* Orientation** | **1.2145** | **0.50059** | **2.4261** | **2836** | **0.015325** | **0.23291** | **2.196** |

In addition to the main effects, we also found a significant interaction between SF Congruency and Orientation (β = −0.93328, 95% CI [-1.5841, −0.28249], t = −2.812, p = 0.004958), as well as between Template SF and Orientation (β = −1.032, 95% CI [-1.6128, −0.45112], t = −3.4837, p = 0.000502). More interestingly, a significant three-way interaction was observed among SF Congruency, Template SF, and Orientation (β = 1.2145, 95% CI [0.23291, 2.196], t = 2.4261, p = 0.015325), suggesting that participants’ accuracy varied not only based on whether the probe had the same or different SF as the template, but also on its orientation (Collishaw & Hole, 2001; Hsiao et al., 2005; Nagayama et al., 1995; Williams et al., 2009). In Figure 2, we present the average accuracy scores in the familiar face matching task for each factor and level.

**Figure 2.**
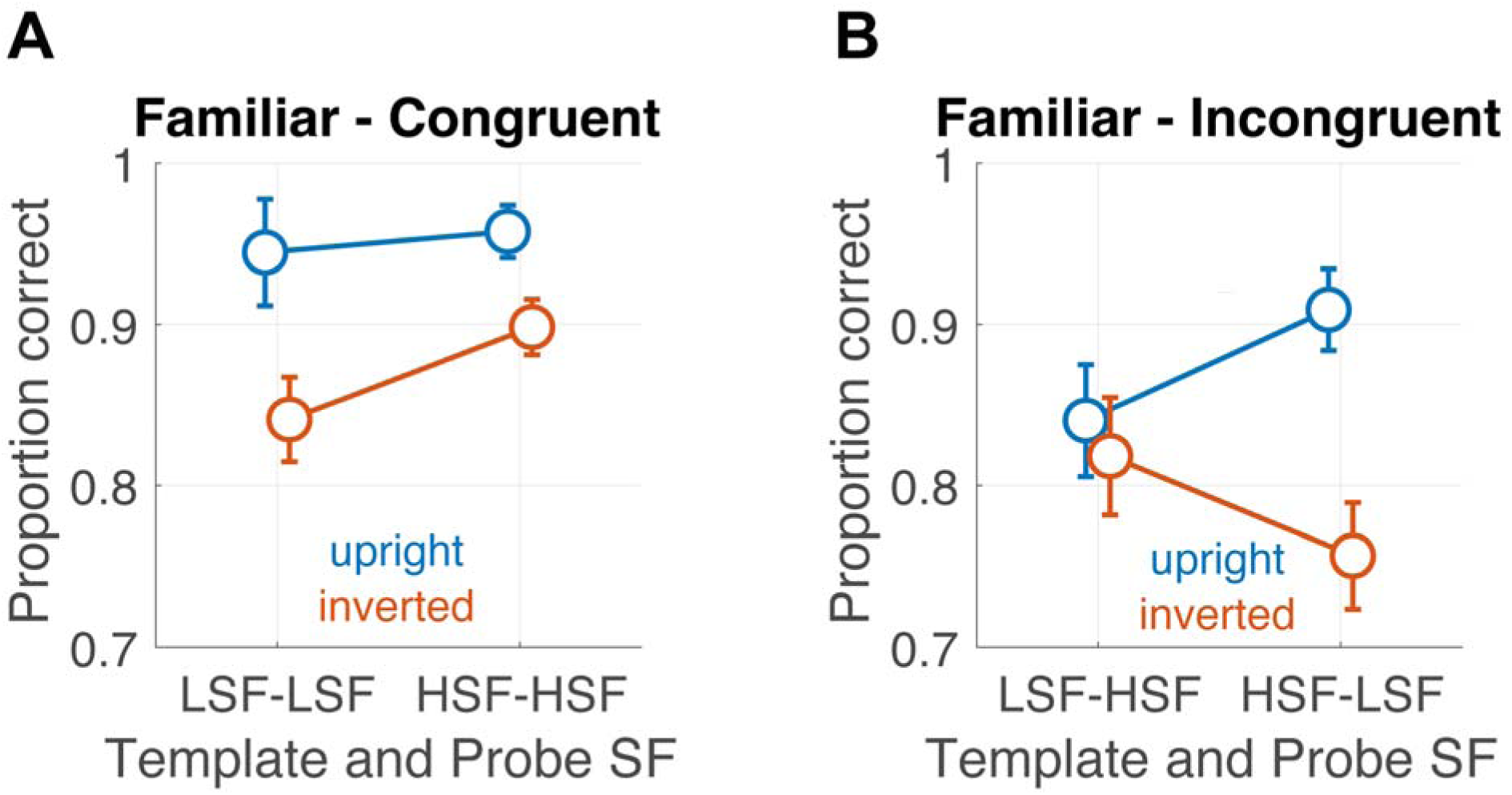
Mean results from Experiment 1 (N = 10). Average accuracy for the familiar face matching task is presented across all conditions. **(A)** The left panel shows the means for the SF congruent condition, where the same SF (either low or high) was used for both the template and the probe. **(B)** The right panel shows the means for the SF incongruent condition, where the complementary SF (either low or high) was used for the probe compared to the template. The data are split by the Orientation factor, with the second face (probe) being -either upright (blue) or inverted (red). Error bars represent one SEM.

### Holistic versus feature-based face processing is engaged depending on probe orientation

To further investigate the three-way interaction, we divided the data based on the SF congruency factor and conducted a separate GLME analysis on each subset of data. For the SF congruent condition (Fig. 2A), the GLME (Eq. 3.1) revealed a main effect of Orientation (β = −1.1478, 95% CI [-1.6724, −0.62322], t = −4.2921, p = 1.89e-05), confirming that performance in the upright trials (M = 0.95, SEM = 0.02) was significantly better than in the inverted ones (M = 0.87, SEM = 0.02). This result suggests that when the probe faces were upright, a holistic processing strategy was used, while participants’ accuracy was driven by a less effective feature-based strategy when matching the upright face of the template with the inverted face of the probe (Itier & Taylor, 2002; Rossion & Gauthier, 2002; Schwaninger & Mast, 2005; Tanaka & Farah, 1993; Taubert et al., 2011). No other main effects or interactions were observed, suggesting that rotation of the probe for the matching task was mostly independent from its spatial frequency (Gaspar et al., 2008; Willenbockel et al., 2010; Young & Burton, 2018).

### Low spatial frequency probes are detrimental to feature-based processing

In the SF incongruent condition (Fig. 2B), the GLME (Eq. 3.2) revealed a main effect of Template SF (β = 0.65797, 95% CI [0.20662, 1.1093], t = 2.8597, p = 0.0043032), and more interestingly, an interaction between Template SF and Orientation (β = −1.0211, 95% CI [-1.5988, −0.44339], t = −3.4672, p = 0.0005417). To follow up on this interaction, we conducted planned contrasts between the four critical conditions. In the upright condition, which involves holistic processing, accuracy was lower when participants saw a LSF template followed by a HSF probe compared to trials in which an HSF template was followed by a LSF probe (LSF-HSFu vs. HSF-LSFu, F(1,1415) = 8.1776, p = 0.004303). In contrast, the opposite effect occurred for inverted probes, which involve feature-based processing. Here, worse performance was observed when participants viewed a HSF template followed by a LSF probe (LSF-HSFi vs. HSF-LSFi, F(1,1415) = 3.9064, p = 0.048295). Crucially, we found that accuracy was higher for upright probes than for inverted ones, but only when the SF template was high (HSF-LSFu vs. HSF-LSFi, F(1,1415) = 31.184, p = 2.81E-08). No significant difference was observed when the SF templates were low (LSF-HSFu vs. LSF-HSFi, p = n.s.).

The findings indicate a differential effect in integrating the spatial content of the probe with that of the template, depending on the probe’s orientation. In the upright condition, participants were more accurate in matching global face information (carried by LSF) to a detailed template (HSF) than vice versa, supporting the idea that upright faces are processed holistically. In this case, HSF templates provided detailed facial representations, facilitating better matches with the global, holistic information in LSF probes. In contrast, inverted probes required normalization of the stimulus (i.e. mental rotation), introducing a perceptual cost, that was particularly high for LSF probes (Schwaninger & Mast, 2005), as also noted, albeit only numerically, in the congruent condition. In these conditions, the perceptual system seems to shift from holistic to feature-based processing, relying on individual features (carried by HSF) rather than the whole face (supported by LSF). Overall, the results support the hypothesis that inverted faces disrupt holistic processing, necessitating a feature-by-feature matching strategy, as it is reflected in the differential use of spatial frequency in the matching task (Itier & Taylor, 2002; Maurer et al., 2002; Rossion, 2008; Rossion & Gauthier, 2002; Tanaka & Farah, 1993; Taubert et al., 2011).

### Analysis of reaction times suggest the lack of speed-accuracy trades off

We continue the analysis by testing if the reaction times were affected by the experimental conditions and highlight possible speed-accuracy trades off. The GLME analysis (Eq. 2, see Table 2) revealed main effects of SF Congruency (β = −0.040829, 95% CI [-0.067012, - 0.014646], t = −3.0581, p = 0.0022561), implying that in the congruent trials (M = 0.84 s; SEM = 0.05) participants were faster than in the incongruent ones (M =0.89; SEM = 0.05). Moreover, a main effect of Orientation (β = 0.074661, 95% CI [0.046939, 0.10238], t = 5.2816, p = 1.4162*10^-7^) suggests that participants were faster in the upright trials (M = 0.84; SEM =0.05) than in the inverted ones (M = 0.90; SEM = 0.05).

**Table 2.** Experiment 1 GLME parameters for reaction times.

| Name | *Beta* | *S.E.* | *t* | *DF* | *p* | 95% *CI*<br>*lower* | 95% *CI*<br>*upper* |
| --- | --- | --- | --- | --- | --- | --- | --- |
| Intercept | 0.85225 | 0.04624 | 18.431 | 2049 | 2.61*10<sup>-70</sup> | 0.76156 | 0.94293 |
| **SF Congruency** | **-0.04083** | **0.013351** | **-3.0581** | **2049** | **0.0022561** | **-0.067012** | **-0.014646** |
| **Template SF** | **0.026864** | **0.013467** | **1.9948** | **2049** | **0.046193** | **0.00045405** | **0.053275** |
| **Orientation** | **0.074661** | **0.014136** | **5.2816** | **2049** | **1.42\*10<sup>-7</sup>** | **0.046939** | **0.10238** |
| SF Congruency * Template SF | -0.03464 | 0.018542 | -1.868 | 2049 | 0.061903 | -0.071 | 0.0017265 |
| SF Congruency * Orientation | -0.00232 | 0.019414 | -0.1195 | 2049 | 0.90489 | -0.040393 | 0.035753 |
| Template SF * Orientation | -0.02903 | 0.020132 | -1.4421 | 2049 | 0.14944 | -0.068514 | 0.01045 |

Furthermore, whereas participants show more accurate responses with HSF Template than with LSF Template, in response times, a main effect of Template SF (β = 0.026864, 95% CI [0.00045405, 0.053275], t = 1.9948, p = 0.046193) indicated that participants were faster when present with LSF (M =0.86; SEM = 0.05) compared to HSF templates (M = 0.87; SEM = 0.05) ones. Taken together, these results suggest a lack of a speed-accuracy trade off, since faster trials were associated with more accurate responses (Fig. 3A and B) (Barragan-Jason et al., 2013; Brandman & Yovel, 2012; Schwaninger & Mast, 2005). No other main effect or interactions were observed.

**Figure 3.**
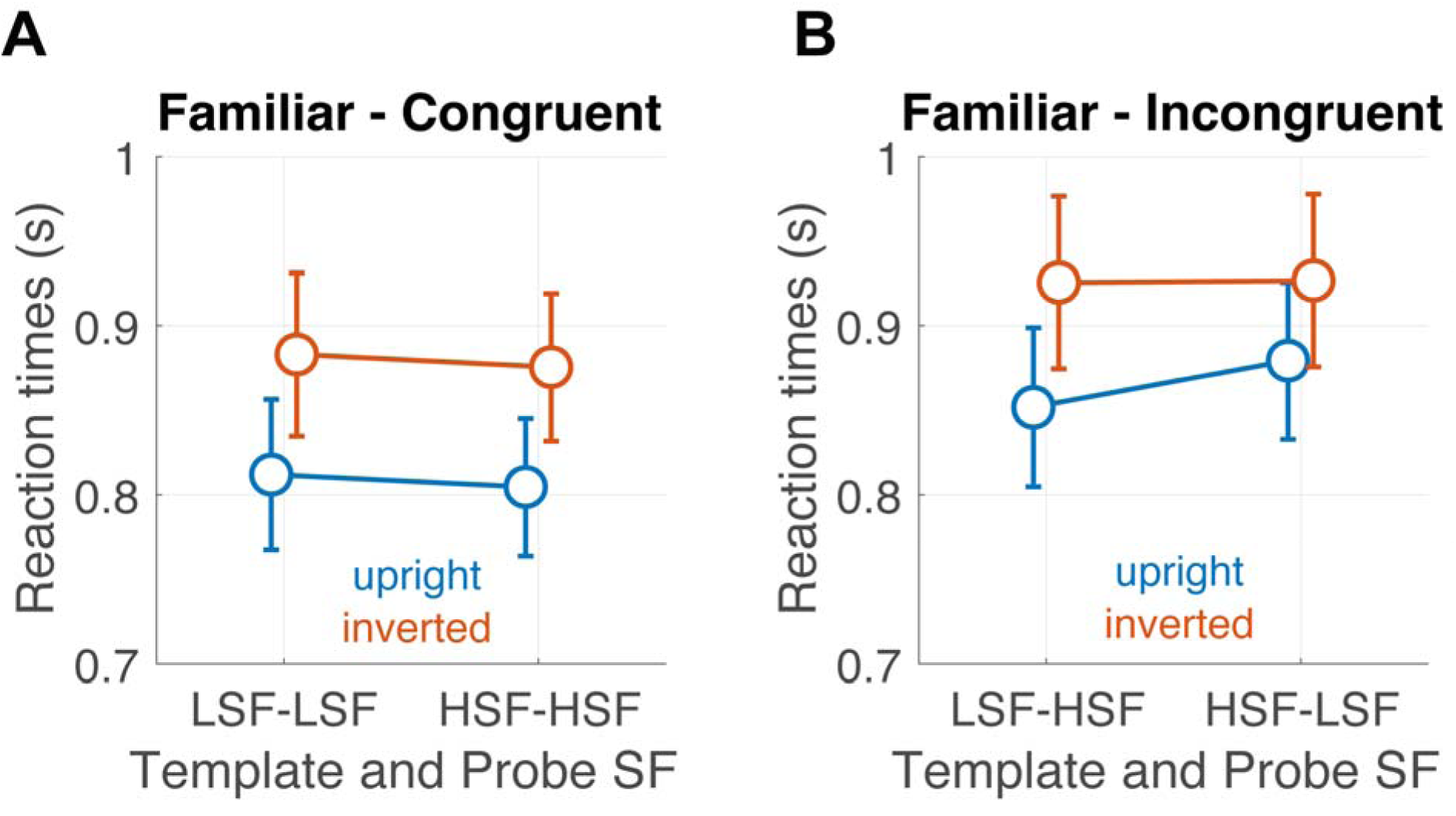
Experiment 1 results (N = 8). Averages RT for the familiar face matching task are presented across the conditions. **(A)** The left panel shows the means for the SF congruent condition, whereby the same SF (either low or high) was used for both the template and match. **(B)** The right panel shows the means for the SF incongruent condition whereby the complementary SF (either low or high) was used for the match compared to the template. The data are split for the Orientation factor, with the second face either upright (blue) or inverted (red). Error bars show one SEM.

## Experiment 2: The effect of familiarity on face integration

### Participants

A convenience sample consisting of twelve participants (8 females and 4 males; mean age 25 years, SD = 3.1909) was recruited to participate in the study. All participants self-reported no history of neurological or visual impairments and had normal or corrected-to-normal vision. They were naïve to the experiment’s purpose and were primarily drawn from university students and staff who participated voluntarily. Written informed consent was obtained, in accordance with the 1964 Declaration of Helsinki. Ethical approval was granted by the local ethics committee at the college of Medical, Veterinary and Life Sciences, University of Glasgow.

### Apparatus, stimuli and procedure

The apparatus and procedure of Experiment 2 were identical to Experiment 1 aside from a few modifications on the stimuli (see Fig. 1). Since the images in the familiar and unfamiliar datasets varied in size, applying a uniform circular mask to all images would have resulted in the loss of essential facial features necessary for recognition. To address this issue, we customized the dimensions of each familiar image, performing specific resizing for each face. This approach allowed us to accurately center each familiar face within the circular mask. To ensure proper image resizing and to address issues related to lighting, we applied filters to reduce the high contrast present in some of the images. Additionally, two images were replaced with those of different celebrities. We applied low and high-pass Gaussian filters with a value of 1.5. The familiarity of participants with the faces of celebrities was assessed through self-report. Each participant was asked to indicate whether they recognized the faces of the presented celebrities, specifying that recognition should not depend on the ability to recall their names but rather on the feeling of familiarity with the face itself. Most participants demonstrated good familiarity with the presented celebrities, with some being able to recognize all the images. However, it emerged that several participants struggled to remember the names associated with the faces, while others did not recognize certain specific celebrities. Additionally, it was observed that some participants knew who certain celebrities were but were unable to correctly identify them from the provided images. The least recognized faces were Uma Thurman (66.67% familiar), Adam Levine (75.00% familiar), and Chris Brown (66.67% familiar). Other than the familiar faces used in Experiment 1, we introduced a new set of unfamiliar faces. These faces were selected from the dataset by Willenbockel et al. (2010), specifically from set 1 used in Experiments 2a and 2b. This dataset includes a variety of expressions, including neutral and happy faces. For this study, we exclusively selected 5 male and 5 female faces with neutral expressions (the same as Experiment 1) to minimize the influence of pose and expression on the results. Each session lasted about 45 minutes, breaks included.

### Experimental design and statistical analysis

The experimental design and statistical analysis were identical to Experiment 1 except that a new experimental condition was added. The full design comprised 32 experimental conditions, determined by the factorial combination of five main factors: *Validity* (validity between template and probe: valid vs. invalid), *Familiarity* (familiar vs. unfamiliar faces), *SF Congruence* (the congruency between the spatial frequency of the template and the probe: congruent vs. incongruent), *Template SF* (the spatial frequency of the template: LSF vs. HSF), and *Orientation* (upright vs. inverted). For the statistical analysis, data were aggregated across the Validity factor. Each of the sixteen conditions was repeated 28 times. Participants performed a total of 448 trials divided in 2 blocks, each consisting of 7 sub-blocks of 32 trials (only two participants performed a total of 512 trials divided into 2 blocks, each consisting of 8 sub-blocks of 32 trials). We collected a total of 5504 trials across all participants in Experiment 2. As for Experiment 1, for each participant and condition (Familiarity by SF Congruency by Template SF by Orientation), we calculated mean accuracy and the mean reaction times (RTs). Mean RTs were calculated only for correct trials. The effects of Familiarity (familiar vs. unfamiliar), *SF Congruency* (congruent vs. incongruent), *Template SF* (LSF vs. HSF), and *Orientation* (upright vs. inverted) on accuracy (0, 1) and reaction times were analyzed using mixed-effects logistic regression (GLME). The models are summarized in Eq. 4 and Eq. 5 in Wilkinson notation (Wilkinson and Rogers, 1973):

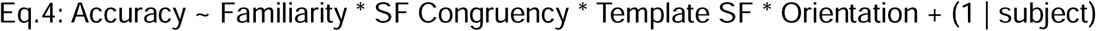

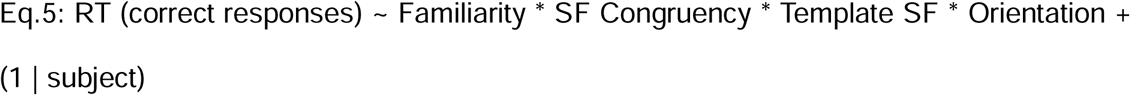

To further investigate the four-way interaction between Familiarity, SF Congruency, Template SF, and Orientation, the data were first divided into the two Familiarity conditions (familiar vs. unfamiliar). Then, for each subset of data, a GLME on accuracy (0, 1) was conducted, examining the effects of SF Congruency, Template SF, and Orientation. From this point on, the analysis follows the steps described in Experiment 1 for the follow-up GLME (Eq. 1 to 3) and t-test comparisons.

## Results

The accuracy in responding ‘equal’ or ‘different’ in the probe matching task was influenced by the factor of interest. The GLME analysis (Eq. 4, see Table 3) revealed significant main effects of Familiarity (β = 0.88364, 95% CI [0.53372, 1.2336], T = 4.9506, p = 7.6242e-07), indicating that familiar trials (M = 0.83, SEM = 0.02) resulted in higher performance compared to unfamiliar trials (M = 0.65, SEM = 0.02). These results confirm that participants successfully recognized the faces of famous individuals and leveraged this familiarity to generate stronger template representations (Castello, Halchenko, et al., 2017; Castello, Wheeler, et al., 2017; Ellis et al., 1978; Klatzky & Forrest, 1984). As shown in Table 3, additional main effects were observed, consistent with Experiment 1. Specifically, performance was better for congruent trials (SF congruency: β = 0.92553, 95% CI [0.57332, 1.2777], T = 5.1515, p = 2.6756*10^-7^) and for upright faces (Orientation: β = −0.43455, 95% CI [-0.74679, −0.12232], T = −2.7284, p = 0.0063853). While several two- and three-way interactions were significant, the most relevant finding for this study — focused on the effect of familiarity, was the four-way interaction between Familiarity, SF congruency, Template SF, and Orientation (β = 1.6183, 95% CI [0.43877, 2.7979], T = 2.6896, p = 0.0071753). This higher-order interaction suggests that familiar faces played a critical role in integrating spatial frequency information across different orientations, particularly when the template SF was congruent with the probe SF. To explore this further, the data were split into two subsets based on the Familiarity factor, and separate GLME analyses were conducted for each subset.

**Table 3.**
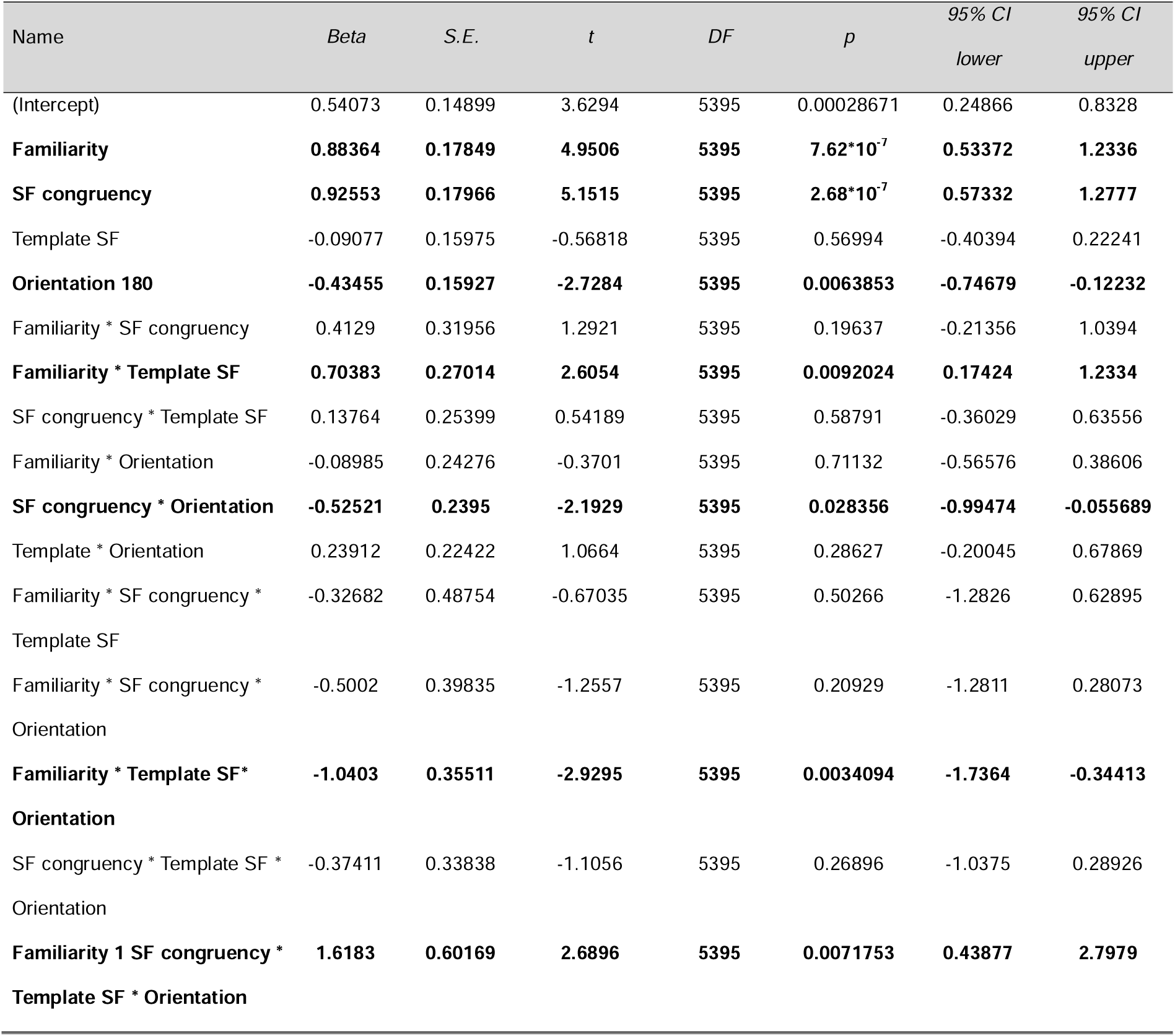
Experiment 2 GLME parameters for accuracy.

| Name | *Beta* | *S.E.* | *t* | *DF* | *p* | 95% *CI*<br>*lower* | 95% *CI*<br>*upper* |
| --- | --- | --- | --- | --- | --- | --- | --- |
| (Intercept) | 0.54073 | 0.14899 | 3.6294 | 5395 | 0.00028671 | 0.24866 | 0.8328 |
| **Familiarity** | **0.88364** | **0.17849** | **4.9506** | **5395** | **$7.62 \times 10^{-7}$** | **0.53372** | **1.2336** |
| **SF congruency** | **0.92553** | **0.17966** | **5.1515** | **5395** | **$2.68 \times 10^{-7}$** | **0.57332** | **1.2777** |
| Template SF | -0.09077 | 0.15975 | -0.56818 | 5395 | 0.56994 | -0.40394 | 0.22241 |
| **Orientation 180** | **-0.43455** | **0.15927** | **-2.7284** | **5395** | **0.0063853** | **-0.74679** | **-0.12232** |
| Familiarity * SF congruency | 0.4129 | 0.31956 | 1.2921 | 5395 | 0.19637 | -0.21356 | 1.0394 |
| **Familiarity \* Template SF** | **0.70383** | **0.27014** | **2.6054** | **5395** | **0.0092024** | **0.17424** | **1.2334** |
| SF congruency * Template SF | 0.13764 | 0.25399 | 0.54189 | 5395 | 0.58791 | -0.36029 | 0.63556 |
| Familiarity * Orientation | -0.08985 | 0.24276 | -0.3701 | 5395 | 0.71132 | -0.56576 | 0.38606 |
| **SF congruency \* Orientation** | **-0.52521** | **0.2395** | **-2.1929** | **5395** | **0.028356** | **-0.99474** | **-0.055689** |
| Template * Orientation | 0.23912 | 0.22422 | 1.0664 | 5395 | 0.28627 | -0.20045 | 0.67869 |
| Familiarity * SF congruency *<br>Template SF | -0.32682 | 0.48754 | -0.67035 | 5395 | 0.50266 | -1.2826 | 0.62895 |
| Familiarity * SF congruency *<br>Orientation | -0.5002 | 0.39835 | -1.2557 | 5395 | 0.20929 | -1.2811 | 0.28073 |
| **Familiarity \* Template SF\***<br>**Orientation** | **-1.0403** | **0.35511** | **-2.9295** | **5395** | **0.0034094** | **-1.7364** | **-0.34413** |
| SF congruency * Template SF *<br>Orientation | -0.37411 | 0.33838 | -1.1056 | 5395 | 0.26896 | -1.0375 | 0.28926 |
| **Familiarity 1 SF congruency \***<br>**Template SF \* Orientation** | **1.6183** | **0.60169** | **2.6896** | **5395** | **0.0071753** | **0.43877** | **2.7979** |

### Familiar Faces Modulate Rotation Processes Depending on SF Congruency

In the Familiar condition, the GLME analysis (Eq. 1) revealed a pattern identical to Experiment 1, with a three-way interaction was observed between SF congruency, Template SF, and Orientation (β = 1.2651, 95% CI [0.28488, 2.2454], T = 2.5307, p = 0.011439). This finding suggests that congruency between the probe and the template was critical for integrating the two faces across different orientations (Fig. 4A). As in Experiment 1, we followed up on the three-way interaction using Eq. 3.1, analyzing the congruent and incongruent data subsets separately. In the congruent condition, we observed only an effect of Orientation (β = 0.45292, 95% CI [-2.0818, −1.0547], T = −5.9907, p = 2.6721*10^9^). This indicates that upright faces were matched more accurately than inverted ones, consistent with the findings of Experiment 1.

**Figure 4.**
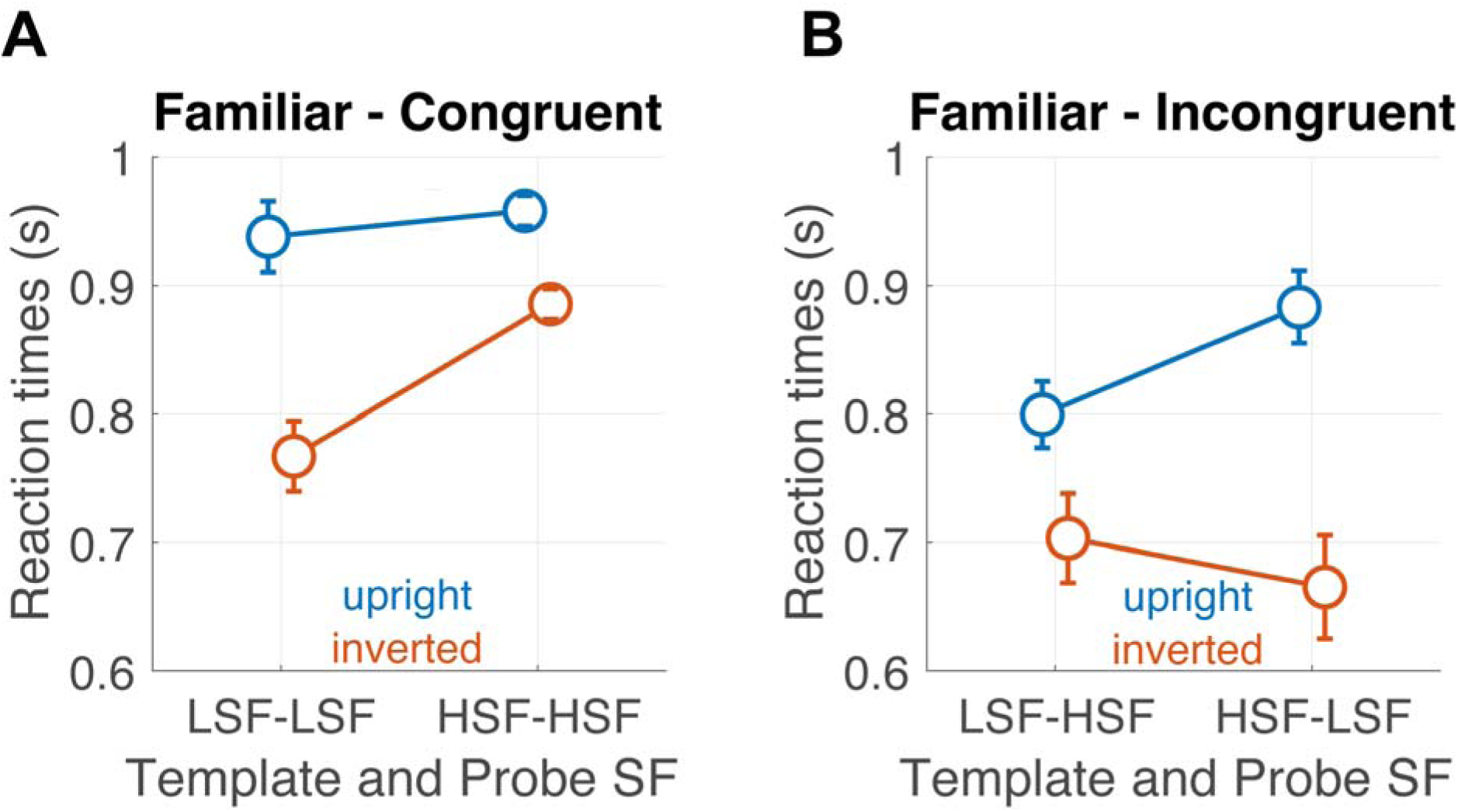
Experiment 2 results (N = 12). Mean accuracy for the familiar face matching task are presented across the conditions. The data are presented with the same convention as Experiment 1.

### Incongruent condition replicates Experiment 1 patterns

More intriguingly, the incongruent condition again exhibited the same pattern observed in Experiment 1 (Fig. 4B), including both main effects and a significant two-way interaction. In the SF incongruent condition, the GLME analysis (Eq. 4) revealed main effects of Template SF (β = 0.62101, 95% CI [0.19084, 1.0512], T = 2.8321, p = 0.0046941) and Orientation (β = −0.53612, 95% CI [-0.89897, −0.17326], T = −2.8985, p = 0.0038111). More notably, an interaction was observed between Template SF and Orientation (β = −0.81541, 95% CI [-1.3601, −0.27067], T = −2.9365, p = 0.003376). To investigate this interaction further, post hoc comparisons were conducted across the four critical conditions. Accuracy for upright probes was higher than for inverted probes when the SF template was high (HSF-LSFu vs. HSF-LSFi: F = 42.495, df = 1341, p = 1.0008*10^-10^) than when it was low (LSF-HSFu vs. LSF-HSFi: F = 8.401, df = 1341, p = 0.0038111). Additionally, in the upright condition, accuracy increased for HSF templates compared LSF templates (HSF-LSFu vs. LSF-HSFu: F = 8.0206, df = 1341, p = 0.0046941). In summary, these results replicate the pattern observed in Experiment 1, providing further evidence for the role of SF congruency and orientation in modulating performance.

### Unfamiliar faces are less susceptible to matching task congruency

The analysis of unfamiliar faces revealed a distinct pattern of results compared to familiar stimuli (Fig. 5). Notably, no main effect of Template SF was observed, indicating that using LSF or HSF templates as input for the matching task did not influence participant accuracy. However, significant main effects were found for SF congruency (β = 0.9177, 95% CI [0.56695, 1.2684], T = 5.1303, p = 3.0978*10^-7^) and Orientation (β = −0.42713, 95% CI [-0.73763, −0.11663], T = −2.6974, p = 0.0070326). These effects were further explained by their interaction (β = −0.52274, 95% CI [-0.99003, −0.055455], T = −2.1935, p = 0.028353).

**Figure 5.**
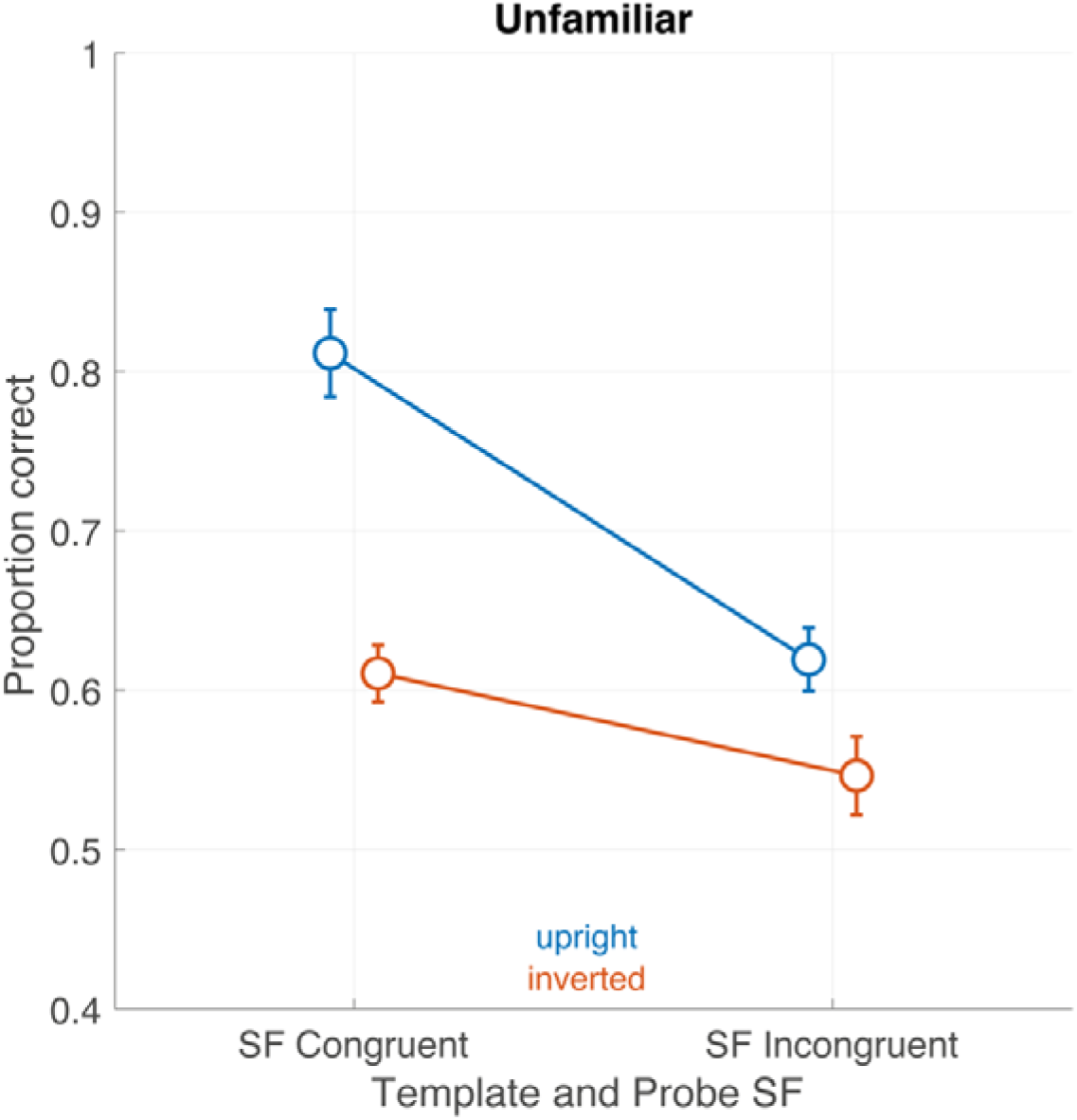
Experiment 2 results (N = 12). Mean accuracy for the unfamiliar face matching task are presented across the conditions. In the x-axis, the data are divided according if the template and the probe had the same (congruent) or different (incongruent) spatial frequency. The data are split for the Orientation factor, with the second face either upright (blue) or inverted (red). Error bars show one SEM.

The data indicate that for upright faces, participants performed better when the spatial frequency of the probe matched the spatial frequency of the template (i.e., the congruent condition), with an advantage of approximately 20% compared to the incongruent condition. This effect was independent of the initial spatial frequency of the template. Conversely, in the inverted probe conditions, performance remained similar between congruent and incongruent SF conditions and was generally lower than in upright probe trials.

### Analysis of reaction times suggests no speed-accuracy trade-off, consistent with Experiment 1

To investigate whether reaction times were influenced by the experimental conditions and to identify potential speed-accuracy trade-offs, we conducted a GLME analysis (Eq. 5, see Table 4). The analysis revealed a main effect of Familiarity (β = −0.038116, 95% CI [-0.073538, −0.0026929], T = −2.1096, p = 0.034955), indicating that familiar trials (M = 0.91, SEM = 0.03) were associated with faster performance compared to unfamiliar trials (M = 0.93, SEM = 0.03)(Ito & Sakurai, 2014; Mulligan & Hirshman, 1995; Yovel & Paller, 2004).

**Table 4.**
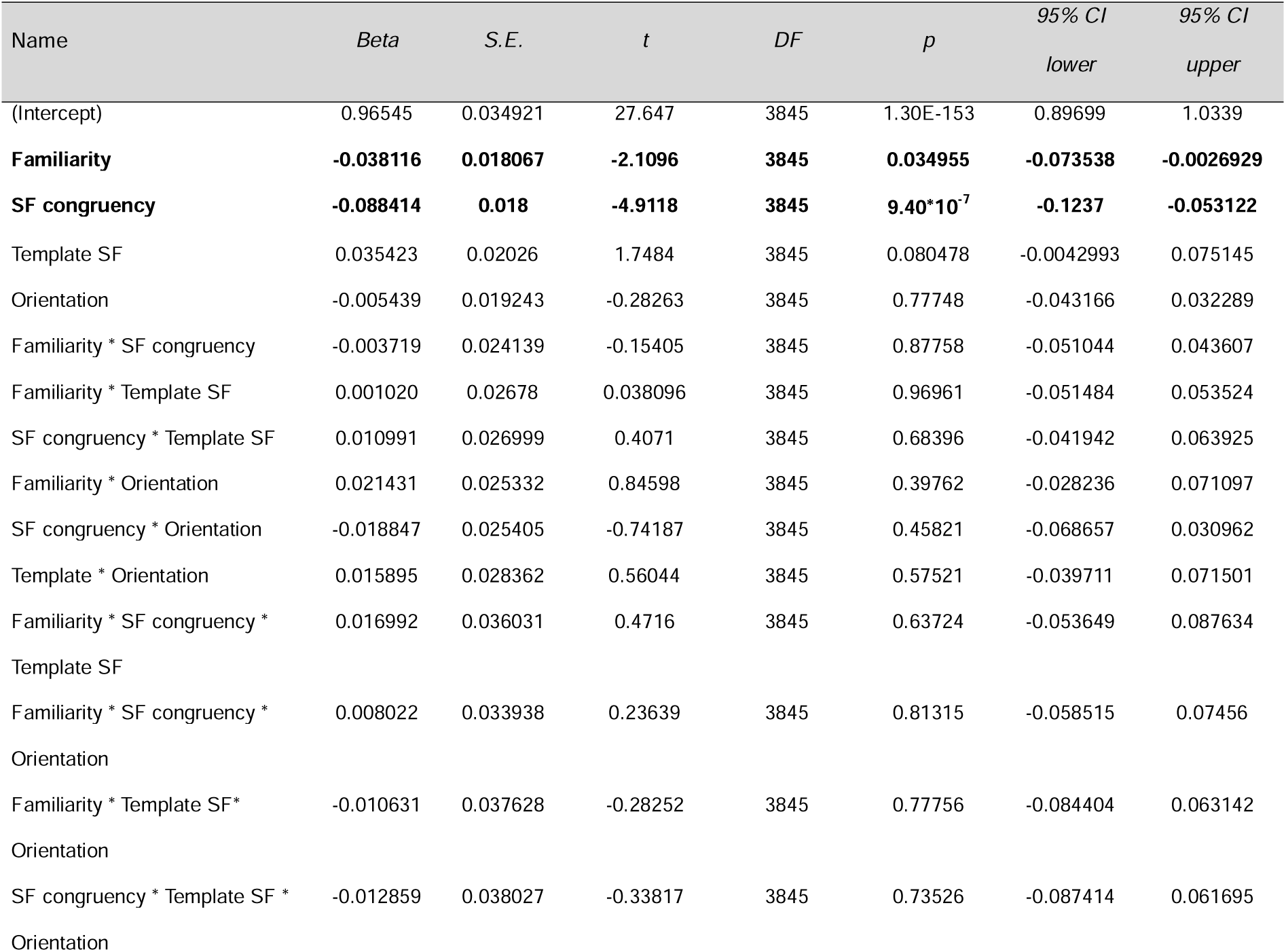
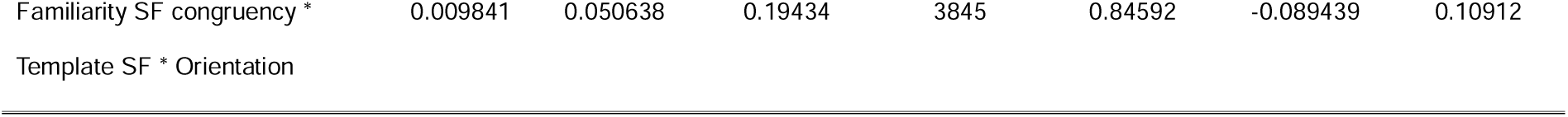
Experiment 2 GLME parameters for reaction times.

We also observed a main effect of SF Congruency (β = −0.088414, 95% CI [-0.1237, − 0.053122], T = −4.9118, p = 9.4019e-07), indicating that participants were faster in congruent trials (M = 0.88 s, SEM = 0.03) compared to incongruent trials (M = 0.97 s, SEM = 0.03). The full set of condition is reported in Figure 6.

**Figure 6.**
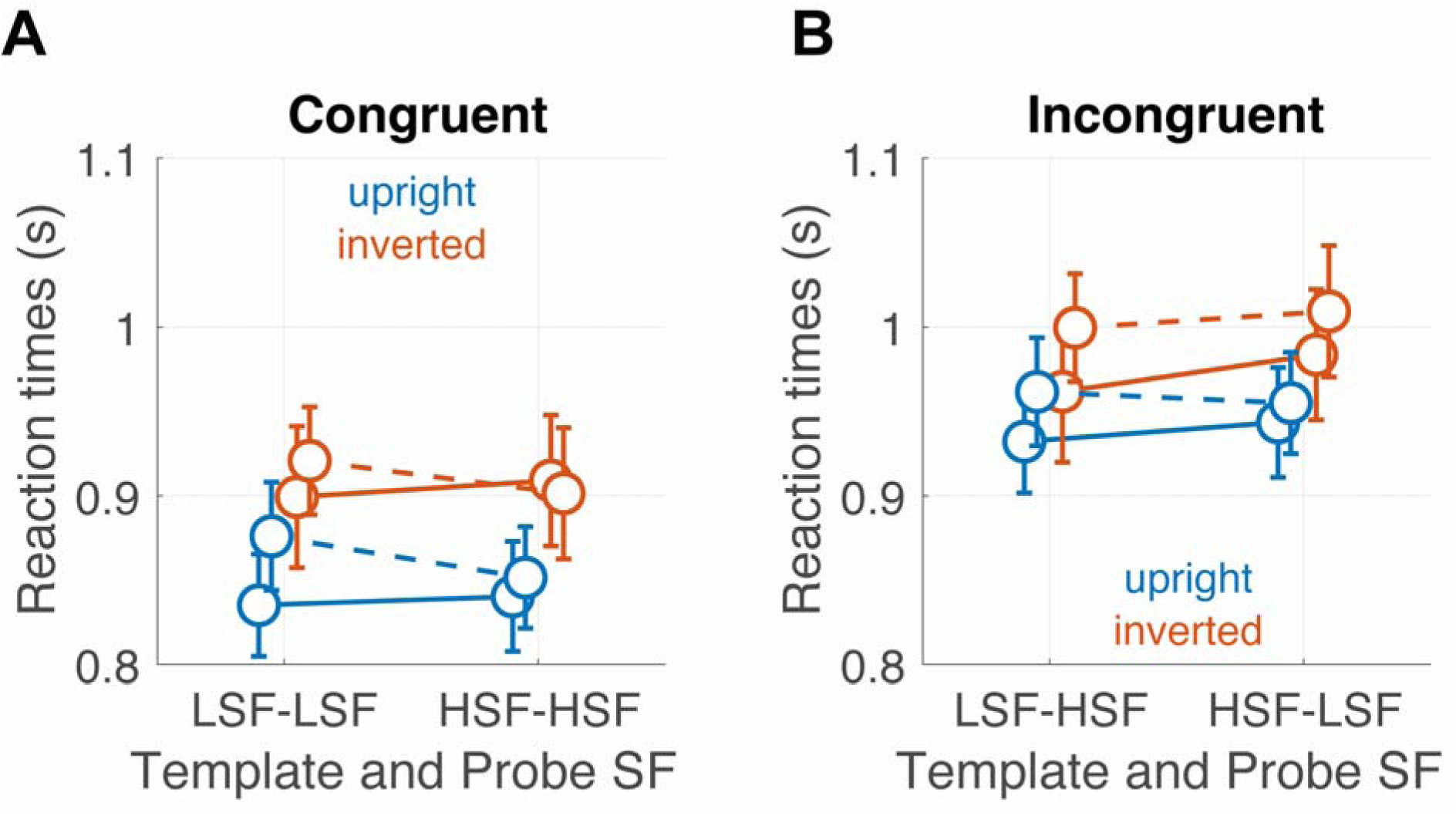
Experiment 2 results (N = 12). Mean reaction times for familiar and unfamiliar faces across all conditions. **(A)** Congruent conditions and **(B)** incongruent conditions. The data follows the same convention as Figure 1. Solid lines represent familiar stimuli, dotted lines represent unfamiliar stimuli.

Overall, these findings support the absence of a speed-accuracy trade-off. The most challenging conditions —namely, incongruent, inverted, and unfamiliar trials, were associated with both slower reaction times and poorer accuracy (Barragan-Jason et al., 2013). Conversely, high accuracy was also linked to faster response times.

## General discussion

This study explored the mechanisms underlying face recognition, emphasizing their flexibility and task dependence in relation to spatial frequency processing (Ruiz-Soler & Beltran, 2006). By examining the ability to match two briefly presented stimuli, we identified patterns in face perception that depend on spatial frequency information, the orientation of the “to-be-matched” face (probe), and face familiarity. These findings align with the so-called Diagnostic Recognition Approach, which posits that the role of spatial frequencies in face perception is determined by the interaction between task demands and the information available in the image (Ruiz-Soler & Beltran, 2006; Schyns, 1998). Overall, our results support the view that complex tasks such as object recognition and categorization cannot be simply predicted by classic visual processing models based on stimulus characteristics alone. Instead, they strongly depend on the interaction between bottom-up sensory input and top-down task-related signals (Bar et al., 2006; Delorme et al., 2004; Lee & Mumford, 2003).

As expected, basic properties of the stimulus, together with task demands, influenced matching accuracy. We observed that in both experiments, congruent trials — where the spatial frequency of the probe matched that of the template, led to higher accuracy. In this case, the strength of the representation provided by the template (whether low or high SF) was less important for matching, as it was overridden by a simple strategy based on stimulus characteristics. This strategy was most effective for upright stimuli but was impaired when the probe required a normalization operation (i.e. mental rotation of the stimulus (Schwaninger & Mast, 2005)). Notably, this effect was observed for both familiar and unfamiliar faces, reinforcing the idea of a predominantly feature-based stimulus-matching strategy. These findings are partially consistent with the results of Williams et al. (2009), who demonstrated that participants exhibited reduced accuracy when verifying and matching complementary faces (i.e., stimuli that do not overlap in any specific combination of spatial frequency and orientation) compared to identical faces. This result highlights that, beyond holistic processing of faces (Richler & Gauthier, 2014; Sergent, 1984; Tanaka & Farah, 1993; Young et al., 1987), a simpler feature-matching strategy is also operating (Audette et al., 2025; Castello, Wheeler, et al., 2017).

We also noticed that in these congruent trials, HSF templates led to higher accuracy than LSF templates (Fiorentini et al., 1982). This finding suggests that providing a detailed template of the face stimuli strengthens the representation for subsequent matching, shaping the interpretation of later conditions. However, this result appears to contrast with some previous literature suggesting that low spatial frequencies might be sufficient for face recognition, with high spatial frequencies being redundant (Ginsburg & Arthur, 1980). Additionally, as discussed below, our results may contribute to the understanding of the coarse-to-fine approach, suggesting that spatial frequency integration is not fixed but dynamically adapts to task contingencies and the specific characteristics of the presented image (Ruiz-Soler & Beltran, 2006; Schyns, 1998)

Among our main findings, we also replicated the well-known inversion effect (Rossion & Gauthier, 2002; Yin, 1969), in which inverted faces reduced accuracy. This result further strengthens research demonstrating that disruptions in either configural processing (the spatial relationships between facial features) or holistic processing can impair recognition (Itier & Taylor, 2002; Maurer et al., 2002; Rossion, 2008; Rossion & Gauthier, 2002; Tanaka & Farah, 1993; Taubert et al., 2011). Within the extensive literature on the inversion effect, some evidence suggests that face inversion may have a more pronounced impact on LSF processing (Collishaw & Hole, 2001; Nagayama et al., 1995; Williams et al., 2009). We also confirmed this pattern, as LSF inverted faces in the congruent condition were the most difficult to mentally rotate and match.

A consistent finding across both experiments was the impact of face orientation on spatial frequency integration in incongruent trials. Upright faces, which are thought to promote holistic processing (Richler & Gauthier, 2014; Sergent, 1984; Tanaka & Farah, 1993; Young et al., 1987), showed improved accuracy when participants were provided with a HSF template to match a subsequent LSF probe. This result supports the notion that HSFs facilitate the construction of a detailed template for holistic recognition, potentially mediated by the ventral pathway for face perception (Collins & Olson, 2014; Schuurmans et al., 2023), which then integrates the LSF information carried by the probe. In contrast, inverted faces disrupted this integration. In this scenario, we suggest that participants switched to a feature-based processing strategy, which relies on local details rather than global configurations (Itier & Taylor, 2002; Maurer et al., 2002; Rossion, 2008; Rossion & Gauthier, 2002; Tanaka & Farah, 1993; Taubert et al., 2011). This shift is likely driven by the normalization of the stimulus involved in the mental rotation task (Schwaninger & Mast, 2005). The differential effects of orientation align with studies emphasizing the importance of configural and holistic processing for upright face recognition (Maurer et al., 2002; Rossion & Gauthier, 2002). Inverted faces, which are processed more like non-face objects (Hsiao et al., 2005), likely disrupt these mechanisms, necessitating reliance on individual facial features. This shift may explain the observed decrease in performance for inverted faces, particularly in conditions requiring LSF integration. Interestingly, the congruent condition showed minimal orientation effects, suggesting that when SFs are aligned between the template and probe, the visual system may rely less on holistic or feature-based strategies and more on direct SF matching (For a review on holistic processing, see Richler & Gauthier (2014). However, see also Audette et al. (2025), who argue that part-based rather than holistic processing predicts individual differences in face recognition abilities).

The results observed in the incongruent condition also align well with a temporal integration framework, according to which spatial frequency information reaches different parts of the cortex at different times, potentially promoting or hindering information integration for the matching task. For example, we might expect that the HSF template travels toward higher cortical areas, such as the inferotemporal cortex, strengthening its signal approximately 150 ms after stimulus presentation and beyond (Goffaux et al., 2011). In contrast, the LSF probe might quickly activate earlier visual areas, but dissipate within 150 ms. These approximate timing estimates align well with our stimulus presentation, in which the probe was presented 100 ms after the template. On the other hand, an LSF template may decay too quickly to establish a strong visual representation for the matching task. By the time the slower HSF signals reach higher-level areas, the opportunity for integration may have passed. We suggest that this hypothesis could be tested by modulating the delay between the template and the probe.

Classical models of vision propose that visual stimuli are initially decomposed into primitive components characterized by their spatial frequency spectrum (Valois & Valois, 1990). Within this framework, the processing of visual information is thought to follow a coarse-to-fine sequence: the visual system first extracts an object’s general shape, conveyed by low spatial frequencies, before progressively integrating finer details carried by high spatial frequencies to form a coherent representation (Bar et al., 2006; Bullier, 2001). Bar et al. (2006) proposed that LSFs are rapidly relayed to the orbitofrontal cortex to generate predictions (Rao & Ballard, 1999), which are then fed back to inferotemporal areas to guide HSF processing and facilitate rapid recognition (Kveraga et al., 2007). This hierarchical process is reflected in anatomical pathways that carry SF information separately from the retina to higher visual cortices (Bullier, 2001; Livingstone & Hubel, 1987; Valois & Valois, 1990). Predictive mechanisms are well-documented in object perception and also influence face processing, even as peripheral information is integrated with the foveal stimulus following eye movements (Buonocore et al., 2020; Huber-Huber et al., 2019).

Our study demonstrates that spatial frequency integration, at least for faces, does not strictly follow this coarse-to-fine approach but is strongly task-dependent (Ruiz-Soler & Beltran, 2006; Schyns, 1998), influenced by stimulus normalization (e.g., rotation) and familiarity. The first key observation is that in our experiments, LSF templates were weak, suggesting that they may not have been optimal for activating a face representation for matching. This was particularly true for unfamiliar faces (see below). In contrast, when familiar faces were used, fine details (HSFs) served as a template for integrating global, holistic information (LSFs), making the effect more pronounced. The absence of this pattern for unfamiliar faces suggests that familiarity enhances template strength and guides SF integration (Tong & Nakayama, 1999). Second, accuracy was particularly low in the incongruent condition when LSFs had to be rotated. This suggests that despite the presence of a strong template provided by the HSFs, and despite being within the correct time window for integration (see above), the LSF information was not sufficient to activate the rapid global representation suggested by the coarse-to-fine theory. Instead, task demands and stimulus normalization primarily drove performance (Schyns & Oliva, 1996). Finally, Experiment 2 further revealed that familiarity significantly modulates SF integration and matching performance. Familiar faces were associated with higher accuracy and faster reaction times than unfamiliar faces, consistent with the idea that familiarity enhances the encoding of facial features and their spatial relationships (Johnston & Edmonds, 2009; Yan et al., 2017). However, see Burton et al. (2015) and Castello et al. (2017) for arguments against a configural processing account of familiar face recognition. Familiar faces benefitted from holistic processing, whereas unfamiliar faces were more likely processed via feature-based strategies. The ability to leverage pre-existing facial representations likely facilitated more efficient spatial frequency integration, particularly under incongruent conditions. It should be noted that recognizing a familiar face involves not only perceptual processing but also access to stored semantic information (Gobbini & Haxby, 2007; Ramon & Gobbini, 2018). Thus, this result is consistent with most recent accounts highlighting the critical role of top-down, semantic-driven facilitation in object visual exploration at both the behavioral (Enge et al., 2023; Federico, Osiurak, Brandimonte, et al., 2021; Federico, Osiurak, Reynaud, et al., 2021; Federico & Brandimonte, 2020) and neural levels (Federico et al., 2023; Federico, Lesourd, et al., 2025; Federico, Osiurak, et al., 2025). For unfamiliar faces, the absence of a clear template SF effect suggests that participants relied more on congruency and orientation cues than on specific SF content. Overall, these findings suggest that SF integration is task-dependent, particularly in incongruent trials, and is modulated by familiarity. These results align with the flexibility hypothesis proposed by Schyns & Oliva (1996), which posits that the visual system dynamically adapts its use of SFs based on task demands and stimulus characteristics.

In conclusion, the findings presented in this study offer new insights into the interplay between holistic and feature-based processing in face recognition. While upright faces benefit from holistic SF integration, inverted faces disrupt this process, shifting reliance toward individual features. Familiarity further amplifies this distinction, enhancing holistic processing for familiar faces while limiting it for unfamiliar ones. These results challenge the assumption that LSFs universally serve as the primary input for face recognition. Instead, they underscore the task- and stimulus-dependent nature of SF integration, where HSFs can play a dominant role in guiding recognition, particularly for upright and familiar faces. Future research is critically needed to explore the neural counterparts of this interplay, shedding light on the underlying mechanisms and their implications for hybrid models of visual perception.

